# Cellular Respiration and Amino Acid Metabolism Is Altered by Dietary Oligosaccharides in *Salmonella* During Epithelial Association

**DOI:** 10.1101/2025.03.09.642302

**Authors:** Claire Shaw, Poyin Chen, Narine Arabyan, Bart C. Weimer

## Abstract

Dietary prebiotic oligosaccharides are common in people’s diets; however, little is known about how different prebiotics alter the enteric epithelium and microbiome. Here we show two structurally different prebiotic oligosaccharides, human milk oligosaccharides (HMO) and mannanoligosaccharides (MOS), alter the metabolism of colonic epithelial cells and Salmonella enterica sv. Typhimurium in ways specific to each prebiotic. Initially, HMO and MOS addition decreased S. Typhimurium association with epithelial cells. However, gene expression analysis revealed significantly induced expression of Specific Pathogenicity Island (SPI) 1 and 2 with HMO treatment opposed to increased fimbriae expression with MOS treatment. MOS treatment induced the expression of genes for amino acid metabolism in both the host cells and in S. Typhimurium, a metabolic shift that was not observed in the HMO treated cells. MOS treatment also altered respiration metabolism in S. Typhimurium to be more closely aligned to those observed in vivo during gut inflammation, which is opposed to colonization-type expression with HMO. Alteration of virulence observed was found to be prebiotic specific and dose dependent, indicating that some dietary substrates likely alter specific pathogens to change their virulence potential in unanticipated ways that lead to multiple outcomes to potentiate or attenuate enteric infections.

## Introduction

Dietary additives that are reported to selectively feed beneficial gut bacteria, termed prebiotics, are commonplace and advertised to the consumers as a panacea for a myriad of health benefits. The US prebiotic market alone currently exceeds $6 Billion and is growing rapidly [1]. Prebiotic oligosaccharides, complex carbohydrates consisting of between three and 10 monosaccharides, are one prebiotic grouping thought to provide beneficial health effects through their roles as metabolic substrates used primarily by probiotic bacteria [2, 3] and through modulation of the intestinal barrier [4–6].

Though classified together under a single label, prebiotic oligosaccharides are structurally diverse and very different in the overall monosaccharide composition. Fructooligosaccharides (FOS), galactooligosaccharides (GOS), and mannanoligosaccharides (MOS) are all commercially available functional oligosaccharides derived from different source material and with diverse structures [7, 8]. Not yet commercialized for broad consumption, but still considered prebiotic oligosaccharides, are human milk oligosaccharides (HMO). HMOs are a combination of structurally diverse oligosaccharides produced by the mother in breast milk and consumed by the infant, subsequently functioning as a selective bacterial carbon source in the large intestine [9]. The structural and functional diversity of prebiotic oligosaccharides increases the complexity around disentangling substrate-microbe-host interactions and their related health outcomes [10]. Though there has been some research into the mechanisms underpinning positive health outcomes, much remains unknown about the effect of structurally diverse prebiotic oligosaccharides on members of the gut microbiota. Much of the research has focused on the beneficial aspects of prebiotics and their use in combination with probiotic bacteria [11, 12]. In contrast, relatively little research has focused on the impact of prebiotics on important enteric pathogens, including *Salmonella enterica* sv. Typhimurium.

*S.* Typhimurium is the most prevalent enteric pathogen in humans and is responsible for over 80 million cases of foodborne illness and 155,000 deaths per year globally [13]. Dietary prebiotics present one path towards the mitigation of this pervasive enteric pathogen, but further investigation is required before clinical application. In previous studies, prebiotics have been employed for the *in vitro* and *in vivo* control of enteric pathogens such as *Escherichia coli*, *Listeria monocytogenes* and *Salmonella enterica* sv. Enteritidis with limited success [14–18]. The modulation of enteric pathogens by prebiotic substrates is in part suggested to be driven via prebiotic-pathogen binding, precluding pathogen binding to host receptors. Certain prebiotics like HMOs and GOS contain glycan structures similar to those found on the gut epithelial cell surface and used by pathogens for host adherence [19]. Enteropathogenic *E. coli* incubated with GOS prior to host introduction showed significantly decreased host association, but GOS was unable to displace already adhered *E.* coli, suggesting prebiotic-pathogen binding prevented initial adherence to host cells [14]. Similar decoy mechanisms have been shown with *Campylobacter jejuni* and *⍺*-1-2-fucosylated glycans, a component of HMO [20]. Prebiotics have also been shown to modulate the immune system irrespective of commensal microbes through interactions with host Toll-Like Receptors (TLR) and G-protein coupled receptors (GPCR) [21, 22]. More generalized protective effects of prebiotics have been attributed to the production of short chain fatty acids (SCFAs), which are small molecules from commensal microbial sources in the gut that support a healthy gut barrier, regulate energy metabolism, and modulate inflammation [23, 24]. The currently understood mode of action for prebiotics suggests the existence of healthy gut microbiota and epithelia is required, indicating that gut dysbiosis impedes prebiotic function. These cumulative findings suggest that prebiotics act as prophylactics, rather than a treatment for enteric infections.

Utilizing a focused system to examine differentiated Caco2 cells during infection with *S.* Typhimurium without any additional microbiota, we showed two structurally different prebiotic oligosaccharides, HMO and commercially available MOS (BioMos®) differentially drive host-pathogen metabolic crosstalk around two major energy-producing routes that altered redox balance in the host-pathogen system and modulated host-pathogen metabolic interactions. This also led to changes in related amino acid metabolism, mitigating the pathogen’s colonization of host cells via prebiotic-specific mechanisms. Pathogen-host association was attenuated by both prebiotic treatments despite induction of *S.* Typhimurium virulence factor expression. Interestingly, the enzymes needed to digest the host glycan were repressed, which may account for the lack of invasion. These observations indicate that a complex relationship exists between prebiotics and gut pathogens. Dietary supplements in this study altered pathogenic activity in previously unexpected ways, illustrating the impact diet can have on enteric pathogens and highlighting the potential for such supplements to exert off target effects with significant impact to host health.

## Results

### Prebiotic pre-treatment of caco2 cells decreases Salmonella association

Pre-treatment of Caco2 cells with structurally different oligosaccharides altered the combined invasion and adhesion activity of *S. enterica* serovar Typhimurium LT2 in a dose-dependent manner (Figure 1). *S.* Typhimurium LT2 was capable of infecting Caco2 gut epithelial cells, evidenced by successful pathogenic activity in the control condition with no prebiotic pre-treatment. Pre-treatment of the Caco2 cells at all tested concentrations with either BioMos® or HMO reduced the association of *S.* Typhimurium LT2 with the differentiated host cells. When added at 0.1% (w/v), BioMos® showed a 59% reduction (p-value < 0.02) in adhesion and invasion while HMO showed a lesser effect at 28% association reduction (non-significant). At 0.5% BioMos® reduced LT2 association by 54% (p-value < 0.05) and HMO by 59% (p-value < 0.05). The max tested concentration, 1% prebiotic, had a 44% reduction (non-significant) in association in the BioMos® treatment while at this same concentration, HMO addition resulted in an 82% decrease in LT2 association (p-value < 0.02). Increased levels of BioMos® decreased efficacy of the pre-treatment while increased HMO levels increased efficacy.

**Figure 1.**
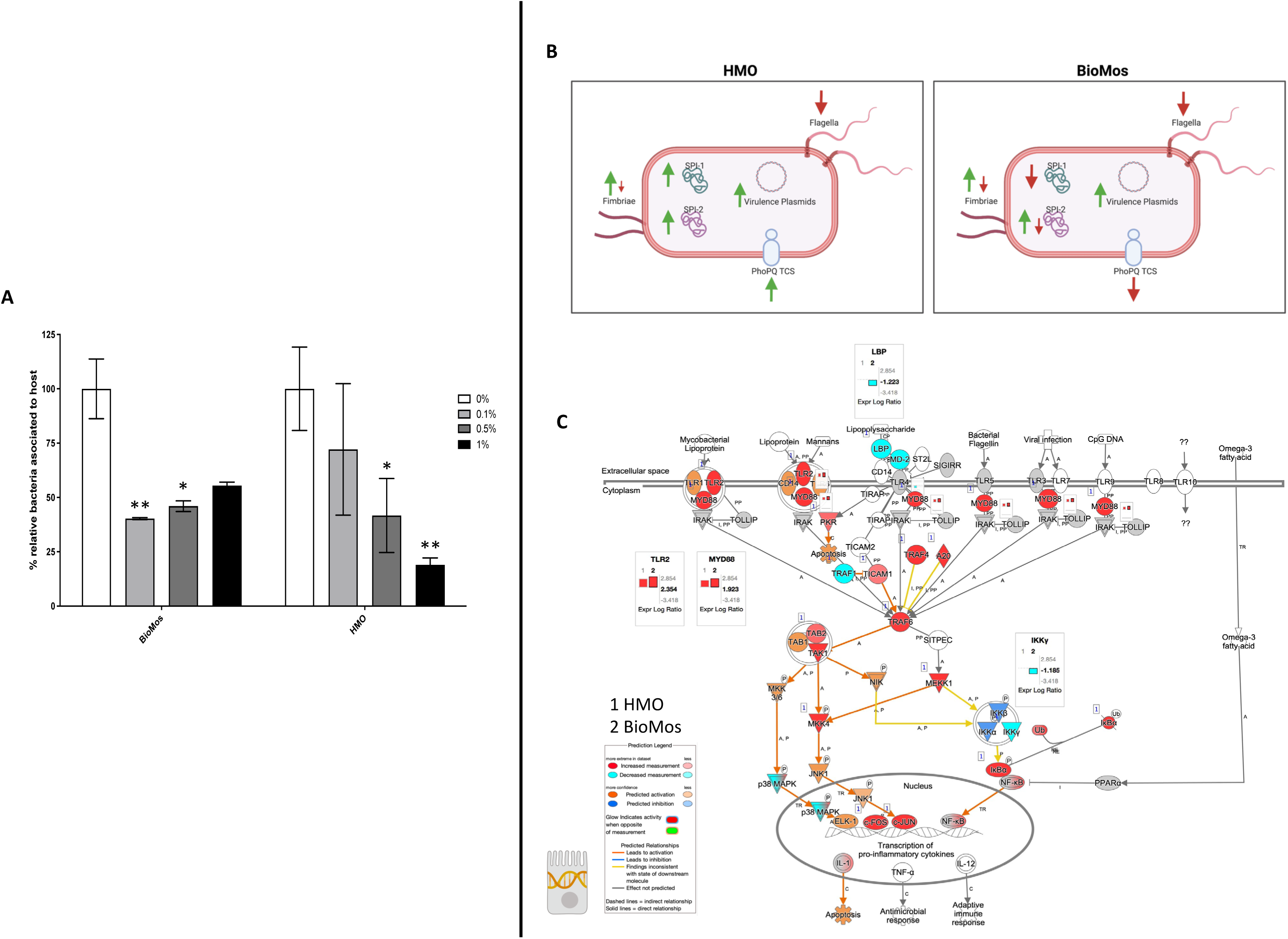
Adhesion and invasion activity of *Salmonella* sp. Typhimurium in response to prebiotic pre-treatment. (A) Prebiotic pre-treatment of Caco2 cells reduces combined invasion and adhesion of *Salmonella* LT2 in an *in-vitro* setting. Prebiotic addition (white bars; 0%) are used as the control for comparison of increasing (0.1, 0.5, and 1%) prebiotic treatment concentrations. BioMos and HMO were added at the noted concentrations to differentiated Caco2 cells for 15 min, followed by co-incubation of *S.* Typhimurium LT2 for 60 mins. Invasion and adhesion were measured with a gentamicin protection assay. * P-value < 0.05, ** P-value < 0.02. (B) Expression of virulence factors in *S.* Typhimurium was evaluated for each prebiotic treatment, using *S.* Typhimurium without prebiotics as the control. Green up arrows represent an upregulation of genes related to each factor while red down arrows represent repression. Specific gene expression information can be found in supplemental Figure 1. (C) TLR gene expression in response to prebiotic treatment was evaluated in the Caco2 cells. The TLR expression pathway from IPA was overlayed with both HMO and BioMos treated Caco2 expression profiles, with no prebiotic Caco2s as the control for both cases. Teal represents decreased measurement for the gene while red represents increased measurement. Blue and orange genes represent predicted inhibition or activation, respectively. Inset graphs display the log_2_ fold-change comparison between HMO (1) and BioMos (2).

*S.* Typhimurium 14028 is 98% genetically identical to *S.* Typhimurium LT2 [25], so the dose dependent response to prebiotic treatment seen in LT2 was predicted to carry over to *S.* Typhimurium 14028 activity. *S.* Typhimurium 14028, being more pathogenic than the lab strain LT2, was used to evaluate the response of host-pathogen interactions to prebiotic treatment via metatranscriptomics and metabolomics.

### Expression of Salmonella virulence factors and Caco2 receptors altered by prebiotics

The reduction in association to host cells of *Salmonella* in both prebiotic conditions led us to investigate if the modulation of known virulence factors in *S.* Typhimurium 14028 and pathogen-sensing Toll-Like Receptors (TLRs) in the Caco2 cells contributed to the defects in adhesion and invasion. Salmonella Pathogenicity Island 1 (SPI-1) and Salmonella Pathogenicity Island 2 (SPI-2) are important gene cassettes that drive *Salmonella* virulence, with SPI-1 supporting initial adhesion to and invasion of host cells and SPI-2 aiding in vacuole escape and systemic spread [26], and together the encode 44 virulence genes.

Thirty-five out of the 44 virulence genes detected in this analysis were induced in the HMO condition, while only 16 of the 44 were induced by BioMos® treatment. Genes related to SPI-1 and SPI-2 were all significantly induced (between -Log_10_ P-value = 6.7 and 17.6, adjP < 3.0e-5) in the HMO treatment but were unchanged or repressed in the BioMos® treatment (Figure 1). SPI-1 genes *sipA*, *sipB, sipC, sipD, prgK, hilA, invA* and *invG* were all unchanged by BioMos® treatment, whereas SPI-2 genes *sseG, sseD, sseC* and *sifA* were repressed and *sseF, sseE, sseB, sifB,* and *sifA* were induced with BioMos® addition (Supplemental Figure 1), suggesting invasion into the epithelial cell was unchanged and escape from the vacuole increased between the treatments.

Fimbriae, structures important for adhesion prior to invasion [27], likewise displayed mixed expression patterns between the prebiotic treatments. *Salmonella.* Typhimurium combined with HMO treated Caco2 cells showed induction of *fimA, fimF, fimY* and *fimZ*. Whereas *S.* Typhimurium combined with BioMos® treated Caco2 cells repressed *fimF* and *fimC,* but significantly induced *fimY* (Log_2_FC = 6.5, -Log_10_ P-value = 11.5), which encodes a mannose-binding type one fimbriae [28]. Expression of virulence genes is in part controlled by two-component systems (TCS), which consist of a membrane-bound receptors and internal response regulators that modulate transcription based on the external feedback. Given their role in virulence, the expression of 13 TCSs across prebiotic treatments was evaluated (Table 1). Generally, TCS-related transcripts in *S.* Typhimurium under the BioMos® treatment were either not found or were repressed. In contrast, HMO treated *S.* typhimurium showed differential expression of all 13 TCSs. Most were repressed compared to expression in *S.* Typhimurium without HMO, though the expression of stress-response TCS, *rpoS*/*rssB* [29], was induced, as was the *arcA* receptor from *arcA/arcB* TCS.

**Table 1.**
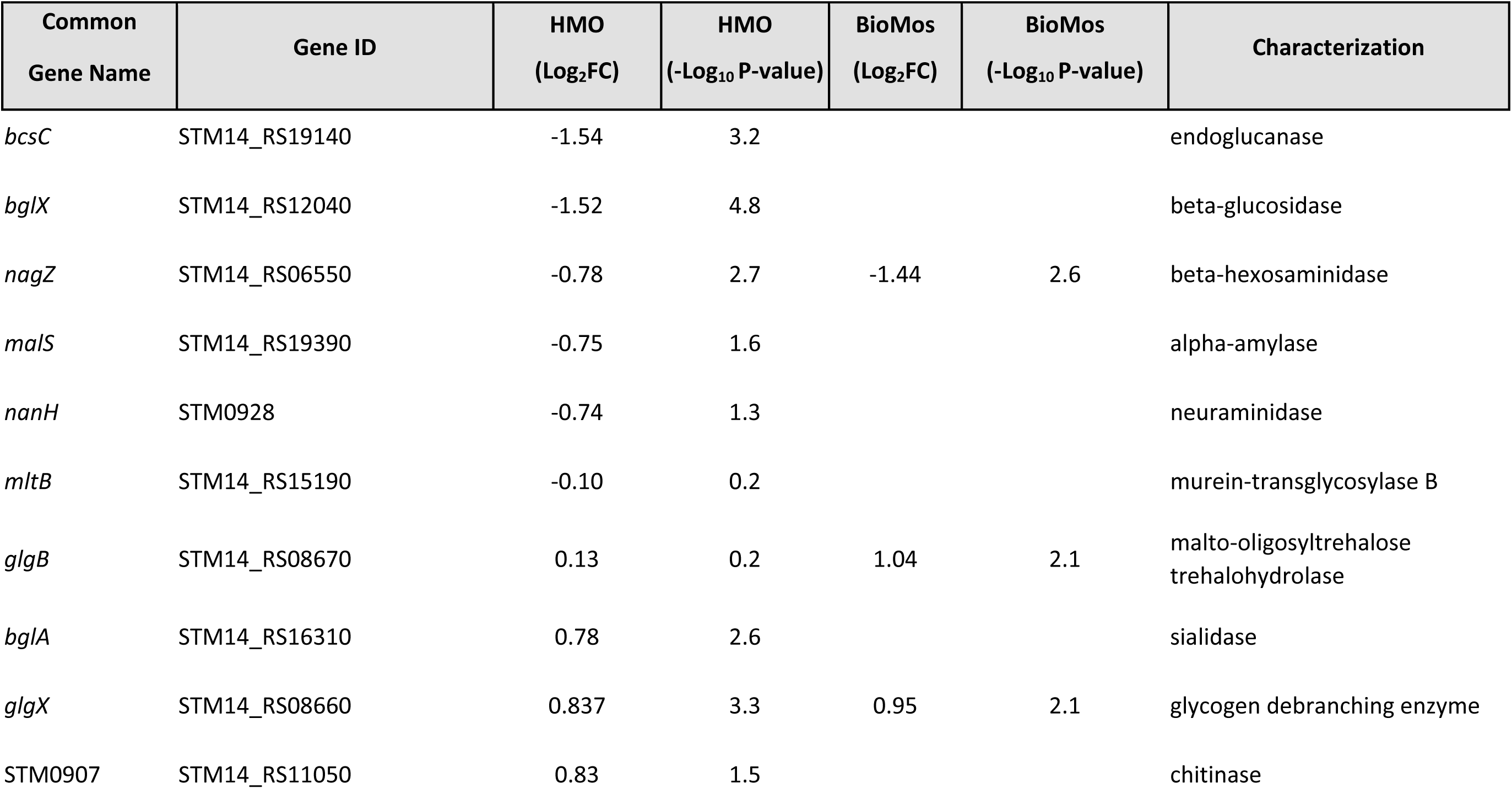
Expression of glycosyl hydrolase enzymes by S. Typhimurium.

| Common<br>Gene Name | Gene ID | HMO<br>(Log<sub>2</sub>FC) | HMO<br>(-Log<sub>10</sub> P-value) | BioMos<br>(Log<sub>2</sub>FC) | BioMos<br>(-Log<sub>10</sub> P-value) | Characterization |
| --- | --- | --- | --- | --- | --- | --- |
| *bcsC* | STM14_RS19140 | -1.54 | 3.2 |  |  | endoglucanase |
| *bglX* | STM14_RS12040 | -1.52 | 4.8 |  |  | beta-glucosidase |
| *nagZ* | STM14_RS06550 | -0.78 | 2.7 | -1.44 | 2.6 | beta-hexosaminidase |
| *malS* | STM14_RS19390 | -0.75 | 1.6 |  |  | alpha-amylase |
| *nanH* | STM0928 | -0.74 | 1.3 |  |  | neuraminidase |
| *mltB* | STM14_RS15190 | -0.10 | 0.2 |  |  | murein-transglycosylase B |
| *glgB* | STM14_RS08670 | 0.13 | 0.2 | 1.04 | 2.1 | malto-oligosyltrehalose<br>trehalohydrolase |
| *bglA* | STM14_RS16310 | 0.78 | 2.6 |  |  | sialidase |
| *glgX* | STM14_RS08660 | 0.837 | 3.3 | 0.95 | 2.1 | glycogen debranching enzyme |
| STM0907 | STM14_RS11050 | 0.83 | 1.5 |  |  | chitinase |

The expression patterns of virulence factors alone did not support the observed prebiotic-driven reduction in host association. We expanded our analyses to include expression of glycosyl hydrolases, which are *Salmonella* enzymes needed to access the host surface and are suspected to be among emerging virulence factors that enable luminal pathogens to achieve contact with the host cell membrane [30] prior to involvement of the T3SS. Glycosyl hydrolases of *S.* Typhimurium under the HMO condition were repressed -*bcsC, bglX, nagZ, malS, nanH, mltB, glgB, bglA, glgX,* and *STM0907* (Table 2). Of the aforementioned genes, BioMos® treated *S.* Typhimurium repressed *nagZ* (Log_2_FC = -1.44, -Log_10_ P-value = 2.6)*, glgX* (Log_2_FC = 0.956, -Log_10_ P-value = 2.1), and increased *glgB* expression (Log_2_FC = 1.04, - Log_10_ P-value = 2.1). These combined observations suggest that HMO treatment led to repression of the glycan degrading enzymes needed to access plasma membrane, thereby hindering bacterial accessibility to the plasma membrane and subsequent host invasion (Figure 1). The importance of these glycosyl hydrolase enzymes in promoting successful invasion of host cells by *Salmonella* was further assessed using *malS* and *nanH* knockouts in *S.* Tyhpimurium LT2 with prebiotic pretreated Caco2 cells, as described previously [30]

**Table 2.** Two-component systems expression among prebiotic treatments.

| TCS Pair | Component | Gene ID | HMO<br>(Log<sub>2</sub>FC) | HMO<br>(-Log<sub>10</sub> P-value) | BioMos<br>(Log<sub>2</sub>FC) | BioMos<br>(-Log<sub>10</sub> P-value) |
| --- | --- | --- | --- | --- | --- | --- |
| ArcB/ArcA | Sensor | *STM14_RS17695* | -0.82 | 2.10 |  |  |
|  | Regulator | *STM14_RS17695* | 2.37 | 12.54 |  |  |
| BaeS/BaeR | Sensor | *STM14_RS22535* | -1.27 | 3.49 |  |  |
|  | Regulator | *STM14_RS22540* | -0.50 | 1.20 |  |  |
| BarA/UvrY | Sensor | *STM14_RS15835* | -0.53 | 1.37 |  |  |
|  | Regulator | *STM14_RS10645* | 1.38 | 4.27 |  |  |
| CpxA/CpxR | Sensor | *STM14_RS21360* | -1.30 | 2.51 |  |  |
|  | Regulator | *STM14_RS21365* | -0.56 | 1.19 |  |  |
| CreC/CreB | Sensor | *STM14_RS24035* | -1.03 | 2.16 |  |  |
|  | Regulator | *STM14_RS24030* | -0.74 | 1.37 |  |  |
| DcuS/DcuR | Sensor | *STM14_RS22600* | -0.04 | 0.06 |  |  |
|  | Regulator | *STM14_RS22595* | 0.92 | 2.38 |  |  |
| EnvZ/OmpR | Sensor | *STM14_RS18570* | -1.07 | 2.79 |  |  |
|  | Regulator | *STM14_RS18575* |  |  |  |  |
| GlrK/GlrR | Sensor | *STM14_RS14045* | -0.91 | 3.09 |  |  |
|  | Regulator | *STM14_RS14035* | -0.93 | 2.05 |  |  |
| KdbD/KdpE | Sensor | *STM14_RS04130* | -1.10 | 1.42 | -1.77 | 3.76 |
|  | Regulator | *STM14_RS04125* | -0.98 | 2.07 | -1.82 | 3.76 |
| NarX/NarL | Sensor | *STM14_RS09720* | -0.58 | 1.20 | 0.85 | 1.76 |
|  | Regulator | *STM14_RS09725* | -0.15 | 0.21 | 1.40 | 1.76 |
| PhoQ/PhoP | Sensor | *STM14_RS06660* | 0.69 | 1.79 | -1.42 | 2.62 |
|  | Regulator | *STM14_RS06665* | 1.26 | 3.95 | -1.46 | 2.61 |
| RstB/RstA | Sensor | *STM14_RS08210* | -1.07 | 2.76 | -1.39 | 2.17 |
|  | Regulator | *STM14_RS08230* | 0.42 | 0.52 | -1.11 | 2.16 |
| TtrS/TtrR | Sensor | *STM14_RS07785* | -1.10 | 2.33 | -1.21 | 2.28 |
|  | Regulator | *STM14_RS07790* | -0.51 | 0.87 | -1.49 | 2.28 |

The importance of glycosyl hydrolase enzymes in supporting *S.* Typhimurium invasion of Caco2 cells was evaluated using *malS* and *nanH* knockouts, which encode for an amylase and sialidase, respectively, in the same prebiotic pre-treated Caco2 cell setup as used for Figure 1. As expected, deletion of *nanH* and *malS* reduced both the adhesion and invasion of *S.* Typhimurium to HMO-pretreated Caco2 cells (Supplemental Figure 2), indicating these sialidase and amylase enzymes are necessary for pathogen access to the host cell membrane. Unexpectedly, BioMos® rescued the invasive phenotype of these same glycosyl hydrolase knockouts (Supplemental Figure 2). This induction of invasive potential indicates BioMos® regulates *S.* Typhimurium virulence mechanisms in ways which promote host invasion, potentially through increased metabolic opportunity for *Salmonella* or through host membrane remodling.

### Host surface receptor expression differentiation with prebiotics

Prebiotic pretreatment of polarized Caco2 monolayers resulted in differential expression of multiple transmembrane receptors in a prebiotic-specific manner, but this differential regulation alone does not explain the difference in the association activity of *S.* Typhimurium (Supplemental Figure 3). To better understand how prebiotics modulated the interaction of the host Caco2 cells and pathogen in this model, we looked at the expression of TLRs, which are integral receptors for initiating an immune response [31]. Both prebiotic pretreatments resulted in the same TLR expression pattern (Supplemental Figure 1). *TLR2* was the only membrane receptor induced with prebiotic pretreatment, and previous work has shown the yeast component zymosan and the HMO structure 3-fucosylactose both induce host *TLR2* expression [32, 33]. *TLR1, 3, 4, 5* and *6* were similarly repressed in both HMO and BioMos® conditions. Expression of the integral TLR signaling protein, MYD88, was induced in both prebiotic conditions, indicating the Caco2 monolayer sensing of *S*. Typhimurium in both treatments (Figure 1).

*TRAF6*, which is downstream of *MYD88* and a regulator of pro-inflammatory cytokine transcription, was also induced in both prebiotic conditions. These data showed prebiotic-specific regulation of transmembrane receptors and immunomodulating TLRs, indicating that host-prebiotic interactions through surface mediated contact regulates intracellular activity. We further investigated the influence of prebiotic-transmembrane receptor interactions on additional extrinsically modulated signaling cascades, i.e., metabolic activity.

### Prebiotic pre-treatment of Caco2 cells modulates the expression of metabolic pathways

The incubation of Caco2 cells with prebiotics prior to infection altered the expression of metabolic pathways, as compared to Caco2 cells infected without any prebiotic pretreatment. Both HMO and BioMos® significantly altered expression of the same 3745 genes as compared to no prebiotic, but BioMos® treatment significantly modulated expression of an additional 3533 unique Caco2 genes compared to HMO treatment (-Log_10_ P > -1.3) (Figure 2). HMO treatment changed the expression of only 191 unique genes when compared to BioMos®. Both HMO and BioMos® treated, then infected, Caco2 cells had significant enrichment of transcripts related to multiple cholesterol biosynthesis pathways, which are a known response to *S.* Typhimurium infection [34], as well as the upregulation of genes related to ketogenesis and oxidative phosphorylation. Overall, HMO and BioMos® treated and *S.* Typhimurium infected Caco2 cells shared enriched expression of the same 11 metabolic pathways, while HMO treated cells had an addition 10 pathways and BioMos® cells had 15 uniquely enriched pathways.

**Figure 2.**
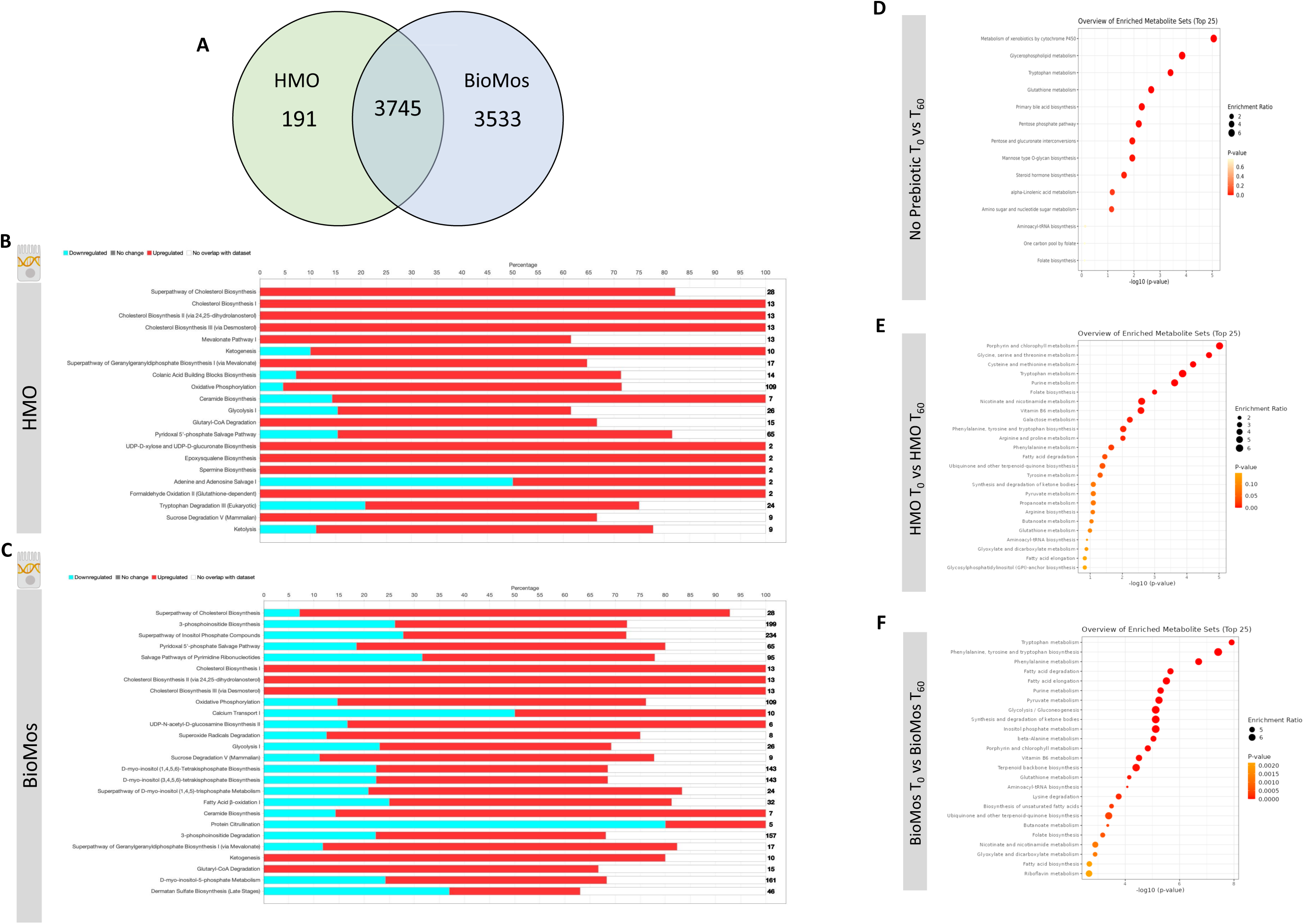
Regulation of gene expression and metabolism of Caco2 cells in response to prebiotic pretreatment. (A) HMO and BioMos treatment of Caco2 cells resulted in 3745 shared significantly expressed genes, 191 additional unique genes in HMO treated cells and 3533 in BioMos treated cells. (B-C) IPA was used to evaluate the expression patterns of the most significant (-Log_10_ P-value > 1.3) metabolic pathways in both prebiotic treatments. Bars represent the percentage of each pathway differentially expressed in each condition, red sections being upregulation, blue downregulation, and white as the percentage of genes found to be unsignificant in the treatment for that pathway. Right hand side numbers represent the total genes in each pathway. (D-F) The metabolic profiles of prebiotic treated Caco2 cells were analyzed using Metabolite Set Enrichment Analysis (MSEA) in MetaboAnalyst. Time 0 with no prebiotic (D), HMO (E), or BioMos (F) was used as the control comparison for the same condition at T_60_. MSEA graph display the enriched pathways at T_60_ for each prebiotic condition, with the dot size representing the portion of the pathway enriched in the dataset and color as P-value.

Caco2 cells pretreated with HMO and infected with *S.* Typhimurium induced expression of epoxysqualene biosynthesis, spermine biosynthesis, and mevalonate pathway I, which are all involved in cholesterol biosynthesis and regulation. Genes related to broad level lipid metabolism are induced in the presence of HMO amid an ongoing infection with *S.* Typhimurium. The differential regulation of spermine, a polyamine derived from arginine and ornithine, in the presence of HMO and *Salmonella* challenge is particularly notable as the production of polyamines by the host has been previously shown to fuel pathogenic activity in *S.* Typhimurium [35, 36]. Additionally, the tryptophan degradation was modulated by HMO treatment but was not significantly altered in BioMos® treated cells. In HMO treated cells, 20% of genes for the tryptophan degradation pathway were repressed, 57% were induced, and 23% were not found in the data. In contrast to the general induction of metabolic pathways seen in HMO treated Caco2s, BioMos® treated cells displayed more mixed regulation, with the majority of significant pathways displaying a combination of repressed and induced genes. Notably the lipid-related ceramide biosynthesis and protein citrullination pathways have significantly altered expression in BioMos® treated cells, though ceramide biosynthesis is primarily induced while protein citrullination was repressed. Treatment of Caco2 cells with prebiotics followed by inoculation of *S.* Typhimurium resulted in the induction of metabolic pathways related to cholesterol metabolism in both treatments, but also resulted in distinct expression profiles of pathways unique to each prebiotic.

Pathway enrichment analysis for metabolites from the control, HMO, and BioMos® treatments compared across 60 mins of prebiotic incubation with polarized Caco2 cells revealed both prebiotic treatments resulted in a greater number of significantly (adj. P-value < 0.05) enriched pathways in Caco2 cells as compared to Caco2 cells without any prebiotic addition (Figure 2). Absent prebiotic pretreatment, metabolism of xenobiotics by cytochrome P450 as the most significantly enriched pathway (-Log_10_ P > 1.3), followed by glycerophospholipid metabolism, tryptophan metabolism, glutathione metabolism and primary bile acid biosynthesis. Both prebiotic metabolic enrichment comparisons also had glutathione metabolism and tryptophan metabolism as an enriched pathway after 60 mins of prebiotic pretreatment/infection (Figure 2). The top five pathways for HMO treated Caco2 cells were porphyrin and chlorophyll metabolism, glycine, serine, and threonine metabolism, cysteine and methionine metabolism, tryptophan metabolism, and purine metabolism. In BioMos® treated Caco2 cells the top five pathways were tryptophan metabolism, phenylalanine, tyrosine, and tryptophan biosynthesis, phenylalanine metabolism, fatty acid degradation, and fatty acid elongation. Notably both prebiotic treatments resulted in the enrichment of multiple pathways related to amino acid metabolism, though not all enriched amino acid pathways in HMO passed the cut-off for significance but still ranked among the top 25 pathways for that treatment. The metabolic profile of HMO treated Caco2 cells without *S.* Typhimurium added supports the gene expression data for HMO treated and infected Caco2 cells, particularly with respect to tryptophan and glutathione metabolism (Figures 2D-F). Though BioMos® treated Caco2 cells also revealed tryptophan metabolites were enriched across 60 mins, tryptophan metabolism did not appear as a top expressed genetic pathway in infected cells treated with the same substrate.

The significant enrichment of different metabolic pathways in Caco2 cells treated with either HMO or BioMos® then infected indicates the prebiotics used in this study exerted an effect on the host cell independent of the presence of any gut flora. Given the observed prebiotic pretreatment-induced differential regulation in Caco2 and the influence of *S.* Typhimurium on diverging metabolic enrichment profiles and gene expression in BioMos® led us to investigate whether such prebiotic-specific effects could be seen for *S.* Typhimurium gene expression in this *in vitro* system.

### Prebiotic treatment of Caco2 cells drives divergent metabolic gene expression in S. Typhimurium 14028

*S.* Typhimurium 14028 was added to the Caco2-prebiotic mixture 15 mins post prebiotic addition and differentially expressed *S*. typhimurium metabolic pathways were determined for each prebiotic in comparison to *S.* Typhimurium with non-treated Caco2 cells. As seen with the Caco2 cells, significant changes in *S.* Typhimurium gene expression were observed in both HMO and BioMos® treatments, but unlike the host cells, BioMos® treatment did not result in the unique expression of any of the bacterial genes (Figure 3). Both prebiotic treatments significantly (adj. P-value < 0.05) altered the expression of the same 2107 genes, although HMO altered an additional unique 2713 genes compared to BioMos®. Visualization of all significantly altered genes resulted in a distinct and opposing expression pattern across the two prebiotic treatments, where most genes that are induced in one treatment are repressed in the other (Figure 3). This observation demonstrates the oppositional effect of the prebiotic treatments on *Salmonella*. The stark difference suggests that bacterial metabolism and virulence will be directly related to the type of prebiotic treatment when consumed in the diet that is not predictable and may lead to potentiation of virulence gene expression but repression of the glycan digestion enzymes that effectively reduces association in a healthy epithelial layer. However, this potentiation for virulence factor expression may lead to unexpected virulence in an inflamed tissue where there is no need to digest the host glycan to gain access to the membrane. The lack of barrier integrity in inflamed tissue synergizes with the metabolic and virulence modulatory effect of prebiotics on *Salmonella* to promote infection and spread.

**Figure 3.**
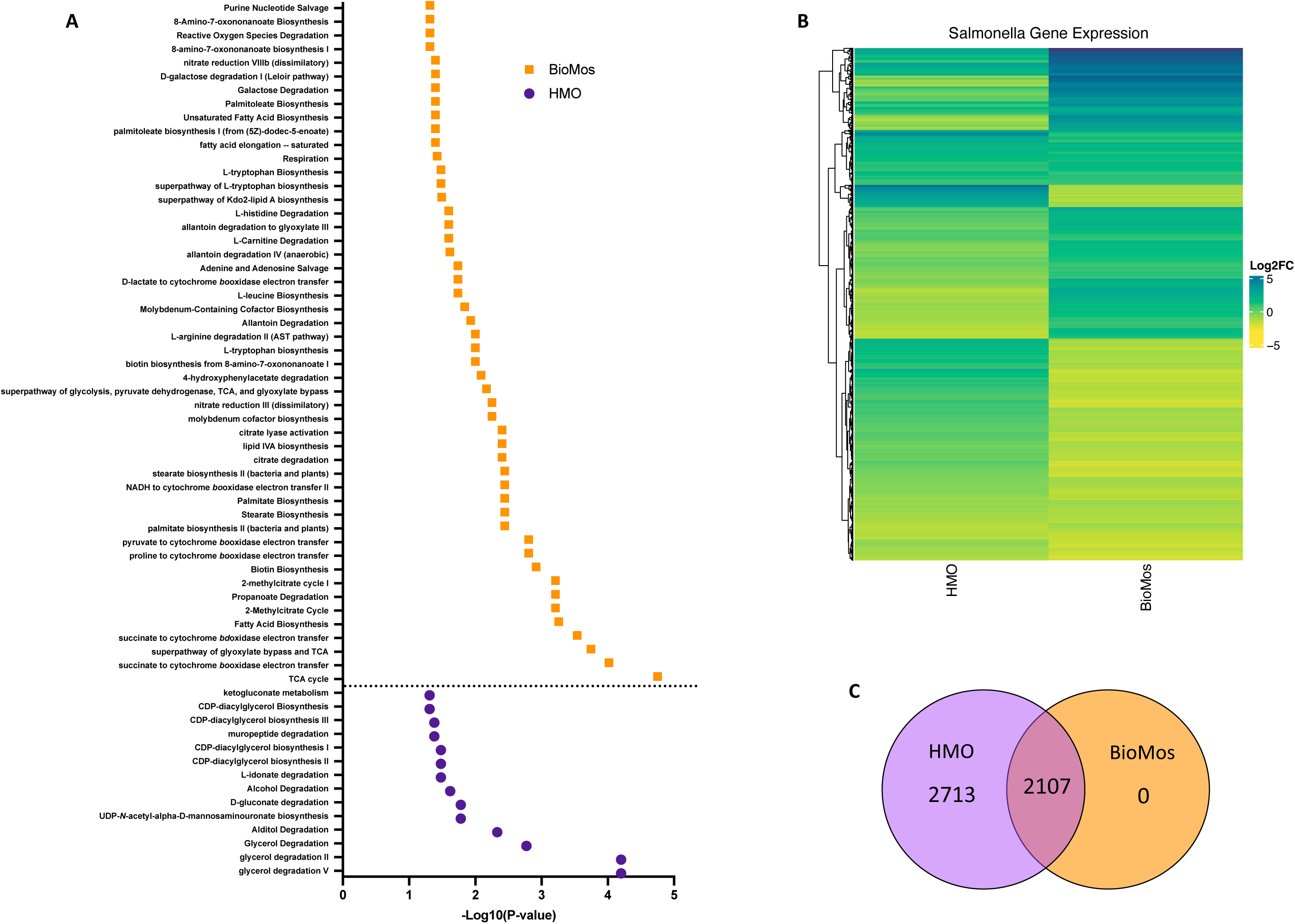
Regulation of gene expression in *S.* Typhimurium in response to addition to Caco2 cells with prebiotic pretreatment. All expression data was collected after 60 minutes of co-incubation with Caco2 cells, with or without prebiotic treatment and in four biological replicates. (A) Enriched metabolic pathway in *S.* Typhimurium added to Caco2 cells pretreated with either BioMos (orange) or HMO (purple). Enrichment determined using BioCyc with a cutoff of –Log_10_ P-value > 1.3 for significance and *S.* Typhimurium added to Caco2 cells and no prebiotic treatment as control condition. (B) Broad evaluation of *S.* Typhimurium gene expression in response to either prebiotic condition using log_2_ fold change data, significance cutoff of –Log_10_ P-value >1.3 and Euclidean distance clustering for genes. (C) 2107 genes were significantly expressed in both the HMO and BioMos treated *S.* Typhimurium, while an additional 2713 unique genes were significantly expressed in the HMO condition and 0 were uniquely expressed in BioMos *S.* Typhimurium.

Targeted analysis of metabolic gene regulation in *S.* Typhimurium identified enrichment of 50 metabolic pathways in the BioMos® treatment and 14 pathways in HMO (Figure 3). HMO and BioMos® treatment did not have any shared enriched pathways in *S.* Typhimurium, revealing distinct reprogramming of metabolism in the pathogen resulting from prebiotic addition. BioMos® treatment resulted in the significant enrichment of multiple amino acid related metabolic pathways including L-tryptophan biosynthesis (-Log_10_ P = 1.31), L-leucine biosynthesis (-Log_10_ P = 1.74), and L-arginine degradation (-Log_10_ P = 2.00). The most significantly enriched pathways in the BioMos® treatment are those related to cellular respiration and energy production. The top four most significant metabolic pathways from the BioMos® treated *S.* Typhimurium were succinate to cytochrome *bd* oxidase electron transfer (-Log_10_ P = 3.54), superpathway of glyoxylate bypass and TCA (-Log_10_ P = 3.74), succinate to cytochrome *bo* oxidase electron transfer (-Log_10_ P = 4.02), and the TCA cycle (-Log_10_ P = 4.75). *S.* Typhimurium from the HMO condition did not show enriched expression for any of these metabolic pathways but rather had enriched expression of genes related to CDP-diaglycerol biosynthesis pathways (-Log_10_ P > 1.31) and multiple glycerol degradation pathways (-Log_10_ P > 1.48). The addition of *S.* Typhimurium to prebiotic pretreated Caco2 cells significantly altered the expression of central metabolic pathways in the pathogen, indicating the addition of these two dietary substrates to host-pathogen systems differentially affects the metabolic communication in host-pathogen metabolic interactions.

### Metabolic profiles reveal distinct shifts in cooperative host-pathogen metabolism after prebiotic treatment

The prebiotic-driven differential expression of genes related to key metabolic pathways in both host and pathogen is supported by the measured metabolites. Whole cell metabolic profiles were determined for each condition (pathogen alone, pathogen and HMO or BioMos®, and HMO or BioMos® alone) between two timepoints, 15 mins post prebiotic addition and immediately after pathogen inoculation (Time 0) and 60 mins post pathogen inoculation (Time 60) with four biological replicates for each combination. K-means clustering with a cluster value of two showed distinct profiles across both time and treatment type (Supplemental Figure 4). Prebiotic type, pathogen inclusion, and sampling time all had marked effects on the metabolic profiles, reflected by the distinct clustering of replicates by treatment combination (Supplemental Figure 5). Caco2 cells with *S.* Typhimurium but without any prebiotic treatment clearly formed two distinct groups. After 15 mins of prebiotic incubation (Time 0) there were distinct clusters by prebiotic type, indicating metabolic profiles of the Caco2 cells were rapidly changed by prebiotic incubation irrespective of pathogen presence. As expected, the metabolic profiles of prebiotic treated and infected samples at Time 0 mirrored that of the corresponding T0 profiles of prebiotic treated uninfected Caco2 cells. More detailed comparison of these HMO vs BioMos® metabolic profiles at Time 0 showed lipids (adj. P-value < 0.02) were increased in the BioMos® treatments as compared to HMO, as were some amino acids like L-kynurenine (adj. P-value = 1.57e-6). The distinct metabolic profiles by prebiotic type observed at Time 0 may be in part due to the addition of exogenous substrates, but combined with the expression data it is clear Caco2 cells alter expression of metabolic pathways in conjunction with prebiotic addition, indicating these noted shifts are not all due to prebiotic addition.

At Time 60 Salmonella presence was the dominant driver of metabolic profile shifts, overshadowing (overtaking?) the effects of prebiotic pretreatment. HMO and BioMos® treated host cells without *S.* Typhimurium formed their own clusters at Time 60, apart from the samples that included *S.* Typhimurium, while Time 60 samples with *S.* Typhimurium formed a cluster set apart from all others regardless of prebiotic treatment type. Though the metabolic profiles of HMO and BioMos® treated cells with *S.* Typhimurium did not cluster separately on the correlation plot by full profiles, a direct comparison of these two treatments showed notable regulation of iron compounds in the BioMos® condition, including precorrin-4 (adj. P-value = 0.02), cobalt-precorrin-2 (adj. P-value = 0.02), FMNH_2_ (adj. P-value = 0.02), and uroporphyrinogen-III (adj. P-value = 0.02). Though not statistically significant but still notable for respiratory pathways, N_2_-succinylglutamate (adj. P-value = 0.2) and pyrimidine-rings (adj. P-value = 0.2) were both among the top 50 features identified in the comparison of BioMos® plus pathogen to HMO plus pathogen at Time 60. The *S.* Typhimurium Time 60 grouping implies a distinctive effect of *Salmonella* on shared metabolism, which overwhelmed the initial prebiotic effect seen for the Time 0 samples, but higher resolution of metabolic products indicated there was still notable differences in the metabolism by prebiotic type.

Across all conditions and time points, 252 metabolites were significant (-Log_10_ P-value > 1.3) and 64 were not significant using the Kruskal Wallis Test. The significant difference for most metabolites (252/316) across all conditions indicates metabolic fluctuation both over time and by treatment (Supplemental Figure 6). All three treatments (*Salmonella* alone, BioMos® and *Salmonella*, HMO and *Salmonella*) at time 60 formed distinctly different clusters and that were also very different from the prebiotics alone at the same time point. Findings from the correlation plot suggest the presence of *Salmonella* distinctly alters the metabolic profile of the combined host-pathogen metabolome, which supports differing expression patterns of the pathogen in either prebiotic treatment. The additional difference between time 60 metabolomes in the BioMos® and HMO treatments without *Salmonella* affirm the observation that Caco2 cells respond to prebiotic treatment and remodel expression of metabolic pathways irrespective of the presence of microbes. The distinct metabolic profiles and large fraction of significantly different metabolites indicate that prebiotic pretreatment of Caco2 cells alters both host and pathogen metabolism.

### Cellular respiration pathways in both host and pathogen are differentially regulated in a prebiotic-dependent manner

The notable metabolic fluctuations and common theme of differentially regulated energy-metabolism related pathways in both the host and pathogen led to the deeper investigation of three major intertwined metabolic routes: the TCA cycle, glycolysis, and oxidative phosphorylation. Caco2 cells displayed different expression patterns for select genes across these three energy-producing metabolic pathways by prebiotic treatment (Figure 4). However, the effect of prebiotic treatment on the expression of genes related to the TCA cycle, glycolysis, and oxidative phosphorylation was much more pronounced in *S.* Typhimurium (Figure 4).

**Figure 4.**
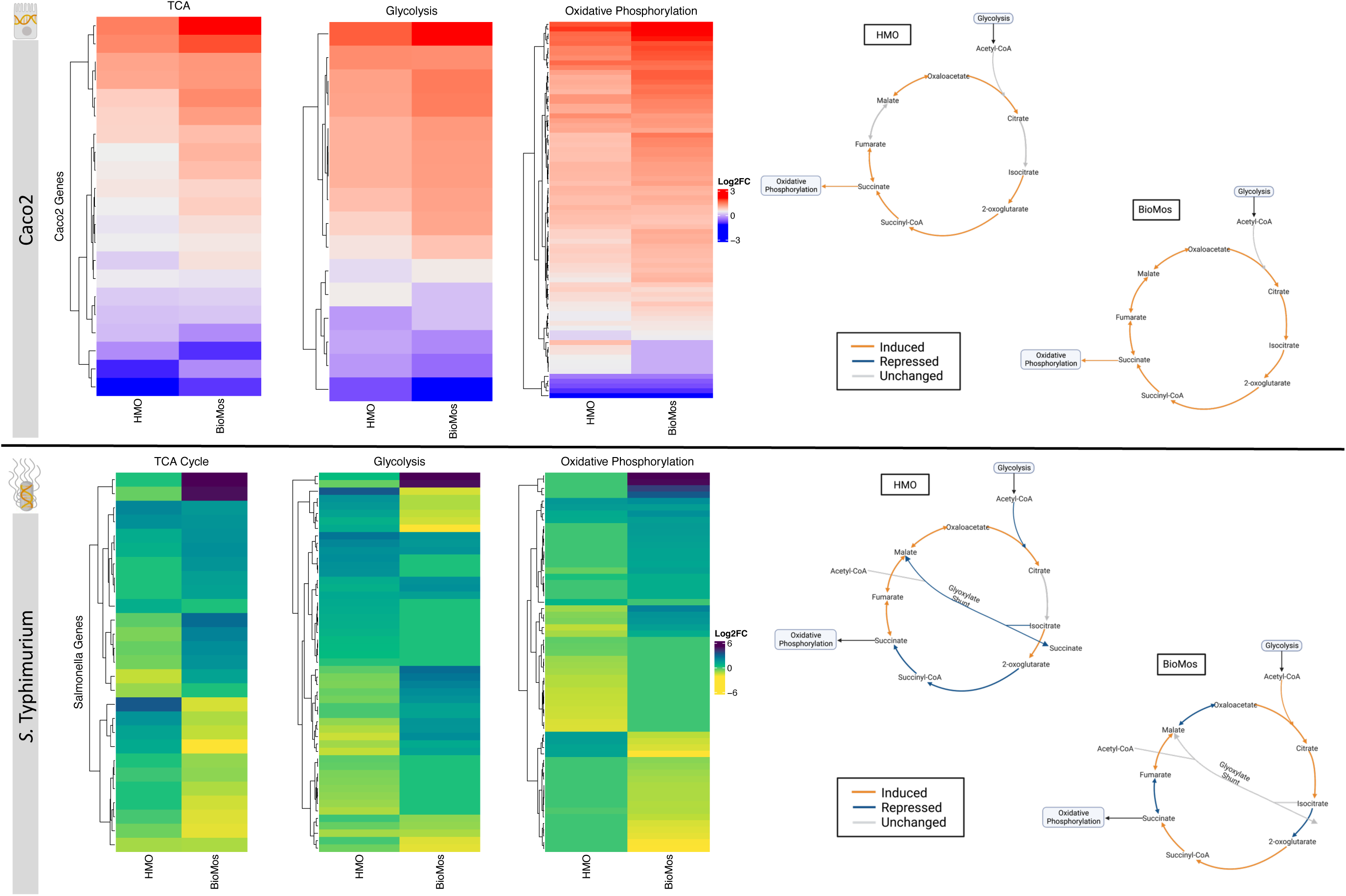
Effect of prebiotic treatment on three major energy-producing metabolic pathways across both infected Caco2 cells and *S.* Typhimurium. Expression of genes related to the TCA cycle, glycolysis and oxidative phosphorylation in repone to prebiotic treatment was evaluated for both Caco2 cells (top) and *S.* Typhimurium (bottom). Caco2 expression of these metabolic respiration-related genes was evaluated using *S.* Typhimurium infected Caco2 cells without any prebiotic treatment added as the control for both HMO and BioMos. *S.* Typhimurium expression was determined using *S.* Typhimurium added to Caco2 cells without any prebiotics as control. All heatmaps used Euclidean distance clustering, Log_2_ fold change expression data, and only significant genes with a cutoff of -Log_10_ P-value > 1.3. Underlying data for the genetic regulation of the TCA cycle (right) was mapped using IPA for the Caco2 cells and BioCyc for *S.* Typhimurium. All genes represented in orange and blue in the diagrams were significantly expressed compared to the control with a cutoff of –Log_10_ P-value > 1.3.

**Figure 5.**
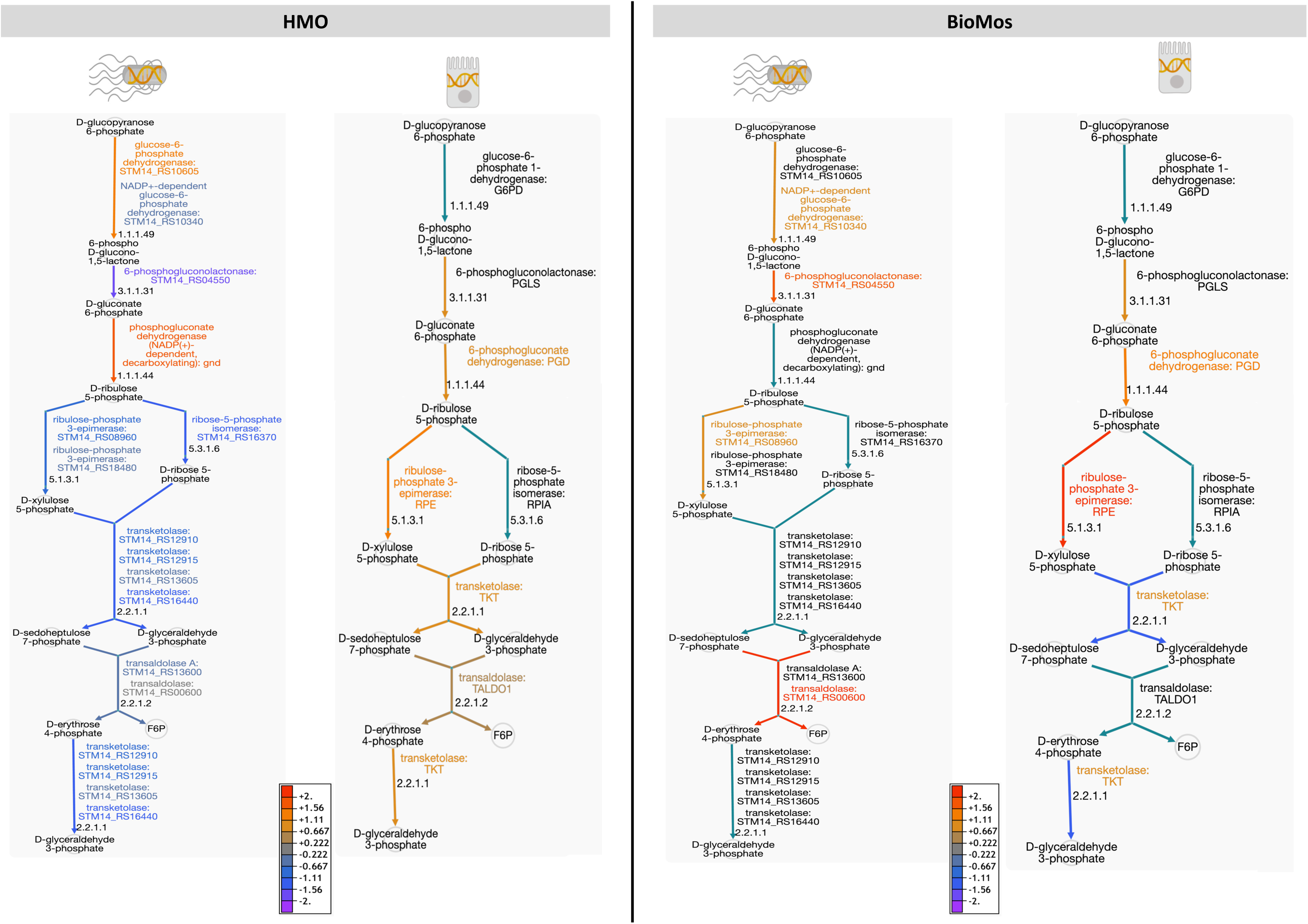
The pentose phosphate pathway is differentially regulated by prebiotic treatment in both Caco2 cells and *S.* Typhimurium. Genetic regulation of the pentose phosphate pathway was evaluated in both *S.* Tyhpimurium (left within each panel) and Caco2 cells (right within each panel) across HMO addition (left) and BioMos addition (right). Expression is represented as log_2_ fold change with no prebiotic add matched host/pathogen cells as control and all regulated genes were significant at –Log_10_ P-value > 1.3. Increased expression is represented by yellow to red and decreased expression by blue to purple. Black text and teal arrows indicate those genes were non-significant or not annotated in the dataset.

**Figure 6.**
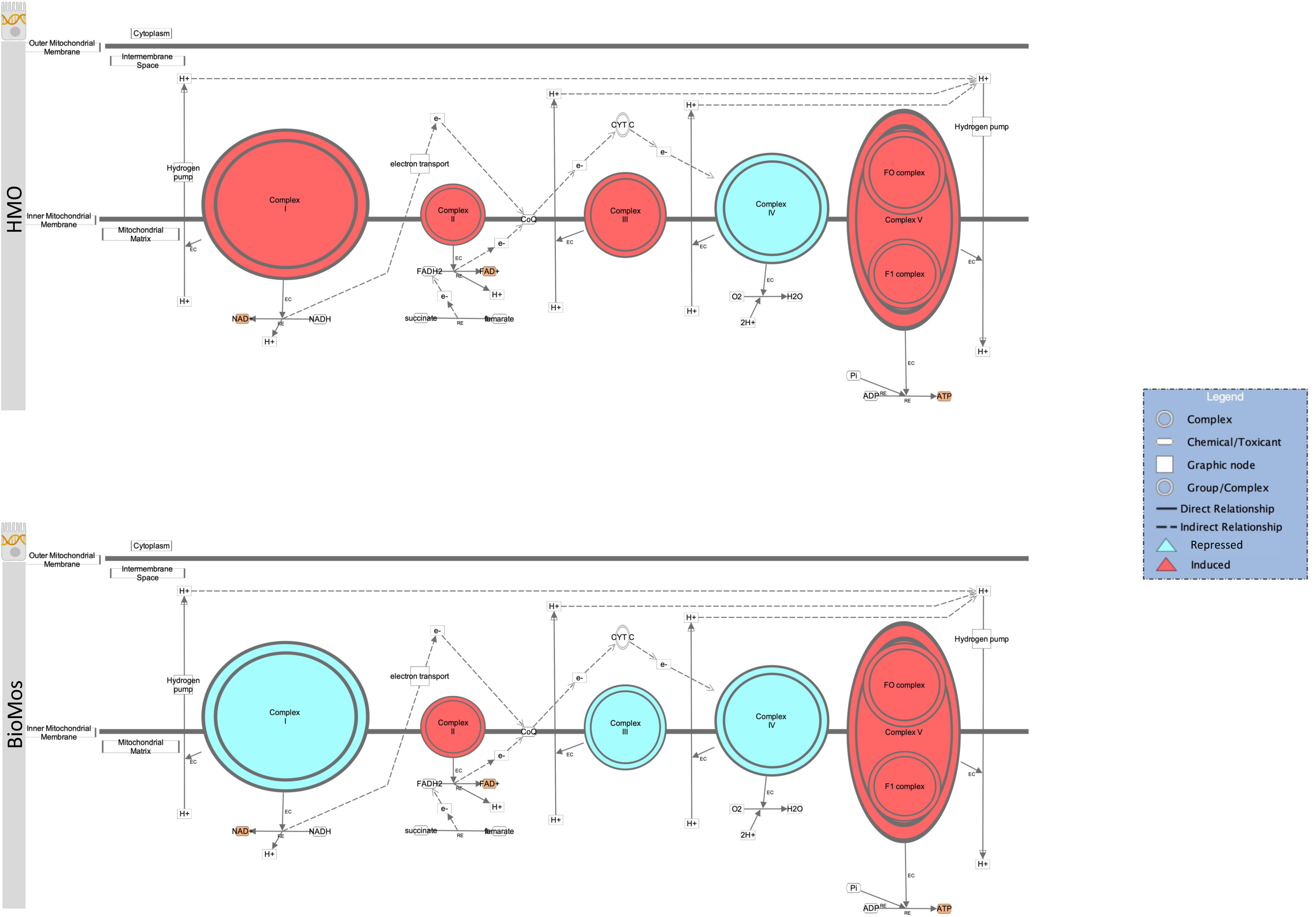
Prebiotic treatment of Caco2 cells challenged with *S.* Typhimurium alters oxidative phosphorylation and expression patterns in the electron transport chain. The canonical oxidative phosphorylation pathway in IPA was overlayed with expression data from Caco2 cells treated with either HMO (top) or BioMos (bottom) and exposed to *S.* Typhimurium for 60 minutes. Red shading in the complexes indicates induction of genes responsible for that complex’s activity and teal shading indicates repression of that complex. Orange shading on the products indicates a predicted increase in concentration due to the expression pattern.

The TCA cycle in Caco2 cells was similarly expressed in both prebiotic treatments, though BioMos® treated Caco2 cells showed an induction of all major reactions in the cycle while HMO treated cells had no differential expression for the steps from citrate to isocitrate and from fumarate to malate compared to untreated cells. In contrast to Caco2 cells, TCA cycle expression in *S.* Typhimurium is different between the two prebiotic conditions (Figure 4). During infection of HMO treated cells, *S*. Typhimurium displayed an incomplete anaerobic TCA cycle where rather than cycling there is a split into oxidative and reductive branches. The repression of the glyoxylate shunt and the repressed reactions from 2-oxoglutarate to succinate seen in the HMO condition indicate TCA regulation in these *S.* Typhimurium cells is following the pattern of early gut introduction and colonization [37]. *S.* Typhimurium infecting BioMos® treated Caco2 cells displayed expression of a more complete TCA cycle, with repression of 3 major reactions (isocitrate to oxoglutarate, succinate to fumarate, and malate to oxaloacetate) that also serve as entry points for external substrates to continue fueling this cycle. This would result in shifting TCA metabolism toward the production of succinate, which is a metabolite with diverse downstream uses including as an electron acceptor to complete the oxidative steps of the bifurcated *S.* Typhimurium TCA cycle [37, 38].

Differential regulation of the TCA cycle, and in particular expression of enzymes related to the oxidative TCA cycle seen in the BioMos® treated cells, is supported by the enriched amino acid metabolism observed in *S.* Typhimurium. *S.* Typhimurium in the BioMos® condition showed induced expression of genes related to biosynthesis of the branched chain amino acid leucine (Figure 3). Arginine and histidine degradation were also enriched in BioMos®, as was tryptophan biosynthesis. All four amino acids, arginine, histidine, leucine, and tryptophan are precursors for substrates in the TCA cycle and their metabolism contributes to TCA cycle activity, either through entry as pyruvate, acetyl-CoA, or *⍺*-ketoglutarate [37]. The induction of genes for a complete complete oxidative TCA cycle in BioMos® treated *S.* Typhimurium in conjunction with the enrichment of amino acid substrates of the TCA cycle suggests complex metabolic regulation by prebiotic treatment and that the regulated pathways do not necessarily confer a host advantage but rather potentiate virulence in the gut environment. Amino acid synthesis in part relies on precursors from another central energy-producing pathway, the pentose phosphate pathway (PPP), so the regulation of this pathway was also examined in more detail.

Two other major energy-producing metabolic pathways that run in parallel, glycolysis and the PPP, display this same pattern of pathway regulation as unique to each prebiotic. As seen with the TCA cycle, Caco2 cells had similar patterns of expression across the prebiotic treatments for genes involved in glycolysis, but *S.* Typhimurium expression was distinctly regulated by treatment. Unlike the TCA cycle and glycolysis, regulation of the PPP in the Caco2 cells differed by prebiotic treatment. The PPP is typically induced in host cells during pathogenesis as a means of creating reactive oxygen species (ROS) from NADPH oxidase ultimately decreasing inflammation through the production of anti-inflammatory cytokines [39]. Likewise, *Salmonella* utilizes the PPP production of NADPH to control redox balance for survival in the host [40]. The finding of differential expression of PPP genes by prebiotic treatment coupled to the known importance of PPP regulation in host-pathogen interactions encouraged a deeper evaluation of the expression patterns of this pathway by treatment and cell type.

Detailed mapping of the pathway revealed repression of the reactions involving transketolase in BioMos® treated Caco2 cells, which drives the latter half of the PPP. PPP repression in BioMos® treated Caco2 cells is not seen in *S.* Typhimurium, which instead induced expression of multiple reactions in the PPP, including transaldolase (*STM14_RS00600,* -Log_10_ P-value = 11.5), which drives production of fructose-6-phopsphate. Contrastingly, HMO treated *S.* Typhimurium displayed repression of the PPP pathway in every step beyond the production of D-ribulose 5-phosphate. Whereas transaldolase was induced in the BioMos® condition, it was repressed in the HMO condition, as was transketolase (*STM14_RS12910*, *STM14_RS12915, STM14_RS16440*) (-Log_10_ P-value > 2.4). While BioMos® induced multiple parts of the oxidative and non-oxidative branches of the PPP in *S.* Typhimurium, HMO generally repressed the non-oxidative branch of the pathway. In Caco2 cells BioMos® repressed the non-oxidative branch and HMO induced this branch, revealing opposing regulation by BioMos® and HMO independent of cell type, host or pathogen.

The final energy producing pathway investigated in this study likewise followed the same trajectory as the two prior paths. Oxidative phosphorylation in both the pathogen and in the Caco2 cells showed distinct modulation of gene expression related to prebiotic treatment. A comprehensive examination of gene expression related to oxidative phosphorylation in Caco2 cells revealed a subset of genes related to complex I of the mitochondrial respiratory chain (*NDUFA3, NDUFA7, NDUFB10, NDUFS6,* and *NDUFS8*) that were repressed in the BioMos® condition (Figure 4). Further, mapping of the respiratory chain gene expression showed repression of Complex IV in both HMO and BioMos® treated cells, but additional repression of Complex I in BioMos®. Notably both prebiotic conditions altered the expression related to the electron transport chain, indicating a level of prebiotic control over this central energy route. Findings in *S.* Typhimurium mirror this observation of electron transport modulation by prebiotics. In *S.* Typhimurium from the BioMos® condition, multiple electron transfer pathways were significantly enriched (Figure 3) whereas these same pathways were not significantly altered by HMO.

## Discussion

Targeted application of probiotics *in vitro* altered gene expression and metabolism of both host and pathogen (Caco2 and S. Typhimurium, respectively) in a prebiotic-specific manner. This observation amends the current paradigm that prebiotics exert protective effects through catabolism by commensal gut microbiota [41] and mitigate infection through competitive exclusion [42]. In contrast to the ligand decoy and/or receptor blocking activity previously described, the prebiotic pretreatment of polarized, colonic epithelial cells performed in this study supported *S*. Typhimurium pathogenesis infection [14, 43]. While prebiotic pretreatment did not abrogate *S.* Typhimurium adhesion to the epithelial cell surface, the rate of invasion and subsequent successful invasion was significantly decreased (p < 0.05).

Together these results indicate that rather than acting as an inert physical blockade, host-protective effects of the dietary prebiotics tested in this study stemmed from a complex orchestration of differential expression of bacterial virulence factors, membrane receptors, and host and pathogen metabolism in a prebiotic-specific manner. Previous work from our lab illustrated the same prebiotics used here remodeled the epithelial cell surface and modulated the expression of receptors, priming the colonic epithelial cells for response to infection with *L. monocytogenes* [15, 30, 44–46]. These previous observations in conjunction with those presented here demonstrate the systemic bioactivity of prebiotic supplements across the gut epithelial barrier, commensals, and enteric pathogens, rather than only in a commensal-specific manner.

Though both were able to reduce host-association at 1% addition, the path by which HMO and BioMos® drove expression changes in both host Caco2 and pathogen cells were prebiotic-specific. The positive correlation between the tested dose and reduced pathogen association in HMO treatment was not reflected in the BioMos® treatment, as increasing BioMos® concentration decreased the host-protective effects. Intriguingly, both prebiotics induced the expression of different virulence-modulating two-component systems (TCS) and other well-documented virulence factors like SPI-1 and SPI-2. Two-component systems are translators for exogenous conditions and allow microbes to tightly regulate and quickly adjust cellular responses to match environmental conditions, both mechanisms key to pathogenic success [47].

All the TCSs examined in this study are important for cellular regulation, but one interesting finding is the repression of membrane receptor CpxA and induction of sensor ArcA in the HMO condition. The CpxAR and ArcBA TCSs cross-regulate each other, with membrane CpxA able to phosphorylate ArcA [48]. ArcA controls multiple genes, including those related to fermentative metabolism and free radical formation, and more interestingly, ArcA activity leads to cell death [48]. In the HMO condition, *S.* Typhimurium shows repression of the membrane component CpxA but induction of the ArcA regulator, potentially indicating *S.* Typhimurium in this condition were pushed towards cell death but ultimately at this timepoint began to repress the external sensor contributing to this cell death pathway [49]. It is possible there was early *S.* Typhimurium cell death with HMO treatment, but over the period of infection this pathogenic attenuation was depressed and overridden through other pathogenic mechanisms.

Despite the induction of multiple virulence factors, regulation of *S.* Typhimurium’s glycosyl hydrolase enzymes and Caco2 receptor remodeling prevented successful membrane access and host invasion. To stage an effective invasion, *S.* Typhimurium must gain access to the colonic epithelial cell membrane, which requires first drilling through a complex mucosal layer made of glycoproteins and oligosaccharides [30]. In a healthy gut this mucosa provides a physical barrier against enteric infection [50], but *S.* Typhimurium possesses multiple glycosyl hydrolases capable of digesting this layer of glycoproteins thus granting the pathogen access to the epithelial cell surface [30]. The reduced expression of these enzymes in the HMO treatment suggests that although key virulence factors like SPI-1 were induced, these cellular mechanisms did not contribute to active pathogenesis in this study as *S.* Typhimurium was not able to physically access to the necessary host receptors.

Interestingly the host was also modulating the associative potential of *S.* Typhimurium and immunological responses via the repression of multiple TLRs, which are expressed in epithelial cells and sense pathogen associated molecular patterns (PAMPs) [31]. TLR2 was the only TLR that was induced, and notably induction was similar in both HMO and BioMos® treatments. Previous work noted that increased TLR2 expression exacerbates *S.* Typhimurium infection through negative regulation of nitric oxide synthase expression and a reduction in epithelial barrier integrity [51]. Expression of TLR-related signaling molecule MYD88 was also induced in both prebiotic conditions. Though this experiment was performed in Caco2 cells, it is notable that previous studies using mesenchymal stems cells, in which *Salmonella* can intracellularly persist and transit through the body [52], found *MYD88* expression was induced in these tri-lineage cells [53]. The induction of *MYD88* in mesenchymal stem cell by the Type III Secretion System (T3SS) resulted in the systemic spread of the pathogen through the host [53], illustrating the exploitation of a host immune response for pathogenic gain and is notable given this same signaling molecule was induced here for Caco2 cells in both prebiotic conditions.

In addition to modulation of cell surface interactions, both prebiotics also regulated metabolic activity in the host and the pathogen. Prebiotics are thought to be privileged for fermentation primarily by commensal bacteria and less accessible to the host and enteric pathogens [54], so the effective prebiotic-driven metabolic control of two non-commensals observed here is an unexpected finding, Metatranscriptome and metabolome analyses indicated energy-producing pathways and related amino acid metabolism were differentially altered in both colonic epithelial cells and *S.* Tyhpimurium in a prebiotic-specific manner . The control of energy-producing redox reactions by *S.* Typhimurium in the gut is important for successful luminal colonization [55] and the diverse metabolic abilities of *Salmonella* allow for adaptation of these pathways to numerous host environments, both extracellular and intracellular [26]. Regulation of these central reactions by prebiotics in this study indicates dietary substrates exert exogenous control of pathogenic metabolism and suggests host diet plays a role in both attenuating and exacerbating enteric infections.

HMO and BioMos® were able to reduce adherence of *S.* Typhimurium to colonic epithelial cells, but this diet-induced reduction was not completely explained by the repression of virulence factor expression or host receptor blocking. The modulation of energetic pathways, and thus energy balance, in both host and pathogen by dietary substrates HMO and BioMos® provides one possible explanation for this attenuated virulence. Predictably, the metabolic composition of samples across time and treatment type (control, prebiotic type, and/or pathogen addition) differed; however, what was notable about these profiles was the distinct clustering of BioMos® treated host and pathogen cells across time. Caco2 metabolites from uninfected BioMos® at T60 clearly clustered together in a group and apart from all other 60 mins samples, regardless of pathogen presence or prebiotic condition, indicating BioMos® distinctly drives Caco2 metabolism divergent from HMO, and *Salmonella* infection overwhelms the metabolic profile in this treatment. This pathogen-driven metabolome in the BioMos® condition suggests that protective effects of BioMos® are transient in the human gut and bacterial infection may be dependent on infectious dose and gut condition. It should be noted that BioMos® is currently used in livestock and primarily studied in the context of animal health, in which BioMos® has been successful for promoting animal health [56]. These findings of less successful mitigation in the human gut context suggests species differences may drive prebiotic efficacy and warrants further research.

This metabolic divergence is further supported by transcriptomic data, which indicate that presence of BioMos® in the media drives metabolic shifts like that seen in inflamed guts [37]; whereas HMO treatment did not show this same effect. The forward push of the TCA cycle reactions, along with the repression of the PPP, and induction of ubiquinone-related metabolism to fuel the mitochondrial electron transport chain in BioMos® treated *S.* Typhimurium suggests BioMos® encouraged aerobic respiration [57] and improved energy production for the pathogen. The TCA cycle in BioMos® treated *S.* Typhimurium displayed a push towards the production of succinate and repression of the downstream production of fumarate, an intriguing finding since succinate is a central compound in driving pathogenesis of multiple enteric organisms [37, 58]. Previous work in *Clostridium difficile* coupled with commensal *Bacteroides thetaiotaomicron* revealed the increased production of succinate by *B. thetaiotaomicron* after antibiotic disturbance supported the proliferation and pathogenesis of *C. difficile* in mice [58]. In this *C. difficile* study, succinate was found to be utilized as a substrate for the regeneration of NAD+ [58] and succinate has likewise been shown to support increased gut colonization and pathogenesis by *S.* Typhimurium [37]. The finding here that BioMos®, but not HMO, drives a more complete oxidative and energetically favorable TCA cycle indicates that prebiotics may alter virulence through the regulation of central energy-regulating metabolic pathways in ways specific to each prebiotic substrate. The control of redox metabolism in conjunction with amino acid metabolism in the gut by *S.* Typhimurium is a key driver of pathogenesis and the switch to aerobic respiration pathways in the BioMos® condition suggests virulence-favoring conditions for *S.* Typhimurium [59].

At the same time, while BioMos® treatment moderated host cell metabolism related to immunological responses, such as reduced calcium transport and expression of multiple lipid-related pathways [60], the addition of *Salmonella* appeared to overwhelm the host protective responses. This takeover may be explained by the prebiotic composition. As BioMos® is a commercial product derived from the cell walls of *Saccharomyces cerevisiae*, it is not pure extracted MOS but instead contains additional substrates derived from the yeast including metal ions and amino acids [56]. *S.* Typhimurium expression showed significant enrichment of amino acid related pathways, which may be a result of these other substrates and not MOS itself, highlighting the off-target effects of commercial prebiotic products may depend on oligosaccharide purity and product composition.

The enrichment of amino acid metabolism in *S.* Typhimurium is notable in the context of the respiration pathway modulation because amino acid-derived compounds, such as fumarate, can act as alternate electron acceptors and contribute to energetically favorable aerobic respiration [61, 62]. Arginine metabolism was also enriched in *Salmonella* in the BioMos® condition, which is of note given arginine serves as a key substrate for proline production and proline is known to modulate oxidative stress in *Salmonella* over the course of infection [63]. The regulation of arginine, leucine, and histidine is also notable given the complete oxidative TCA cycle observed in the BioMos® treated cells, but not in the HMO treated ones. Amino acid regulation is a key component of nutritional immunity between hosts and pathogens [64]. The enrichment of amino acid metabolic pathways able to fuel the TCA cycle through the oxidative branch and towards more efficient ATP production in the BioMos® treatment is a notable finding, as this appears to support pathogenic activity rather than attenuate it.

L-tryptophan, an essential amino acid and central substrate for many bioactive molecules and a key precursor for nicotinamide adenine dinucleotide (NAD+), was regulated in both host and pathogen by BioMos® treatment [65–67]. L-tryptophan biosynthesis was upregulated in BioMos® treated *Salmonella* and generalized tryptophan metabolism was a significantly enriched metabolite pathway in Caco2 cells. L-kynurenine, a direct production of tryptophan degradation, was an enriched metabolite in BioMos® treatment as compared to HMO. L-kynurenine is known to be a neuroactive metabolite with broad neurological effects, both protective and detrimental, in humans [68]. This additional finding of tryptophan regulation by both host and pathogen in conjunction with the more specific measurements of metabolites from this metabolic pathway gives credence to prebiotics potentiating health effects beyond the gut barrier, though more research is needed on these potential systemic effects. Additionally, tryptophan’s pivotal role as a key substrate for redox coenzyme NAD+ and as a precursor to bioactive molecules affecting both host and pathogen makes the enrichment of tryptophan metabolism in *S.* Typhimurium with BioMos® noteworthy and something to consider when testing prebiotic efficacy in pathogenic contexts.

HMO treatment did not elicit the same enrichment of amino acid metabolism in *Salmonella*, likely because the HMO used in this study was a purified non-commercial product with minimal if any additional substrates beyond the complex oligosaccharides. HMO treatment better prevented *S.* Typhimurium association with host epithelial cells and primarily resulted in anaerobic respiration in the pathogen. Somewhat confoundingly expression data for *S.* Typhimurium showed more uniquely expressed genes in HMO (2713) and no uniquely expressed genes in BioMos®. The complexity of the HMO oligosaccharides in contrast with the simpler MOS structure and accompanying yeast cell wall parts may provide an explanation for this finding. Indeed, HMO treatment did not result in the enrichment of many metabolic pathways for *Salmonella* and alterations in host cell expression were minimal. The repression of energy production pathways in *Salmonella* with HMO treatment, known host cell surface remodeling by HMO [15], along with previously understood positive immunomodulatory functions of this prebiotic [69] supports that HMO in healthy gut condition may attenuate enteric infection by *Salmonella.* Pathogenic attenuation by dietary substrates appears highly substrate and host specific, as illustrated by one clinical trial applying FOS to patients with Crohn’s Disease, which produced no clinical benefit to participants [70]. This previous work provides some *in vivo* evidence that prebiotics in an inflamed gut may not always produce the expected ameliorative properties.

Observations from this work indicate dietary substrates may on the surface attenuate virulence but underneath can drive expression of pathogenic-related genes and metabolic pathways, that potentiates virulence in compromised tissues, which is a cautionary finding for the clinical application of dietary interventions. The additional finding here that substrate type influences both the host and pathogenic response points to the complexity of utilizing dietary interventions for the control of pathogens. These results indicate the beneficial effects of dietary prebiotics may be contingent on their addition to an already healthy gut environment, and rescue of a dysbiotic or inflamed gut from infection is specific to only select oligosaccharides. In the context of pathogens, the modulation of invasion by the prebiotic oligosaccharides seen in this study shows promise for the use of dietary substrates as prophylactics, but the diverging effects on pathogenic energy metabolism, which may increase virulent traits, supports the ongoing need for substrate-specific studies across both healthy and dysbiotic gut environments.

## Methods

A graphical representation of the experimental set up can be found in Supplemental Figure 7.

### Oligosaccharides

HMO was isolated and given as a gift to the Weimer lab by Dr. Daniela Barile (UC Davis, CA, USA) [71]. BioMos® is a yeast-derived and commercially available product from Alltech Inc. (Nicholasville, KY, USA). Brief descriptions of each oligosaccharide mixture can be found in Table 3. Both oligosaccharides were made into a 1% working concentration in high glucose DMEM (HyClone Laboratories, Logan, UT, USA).

**Table 3.** Source and content of oligosaccharides used in this experiment.

| Prebiotic | Main Oligosaccharide Structure(s) | Prebiotic Source |
| --- | --- | --- |
| BioMos® | Mannan-oligosaccharides | Proprietary *Saccharomyces cerevisiae* derived product |
| HMO | Five monosaccharides (Glucose, Galactose, N-acetylglucosamine, Fucose, Sialic Acid) combined into complex oligosaccharides | Extraction from human breast milk |

### Bacterial Strain and Growth Conditions

Bacterial cells were grown as previously described [30, 45, 72–74]. Briefly, *Salmonella enterica* subsp. *enterica* serovar Typhimurium strain LT2 was grown for the prebiotic adhesion and invasion panel and *Salmonella enterica* subsp. *enterica* serovar Typhimurium strain 14028s was used for an additional adhesion and invasion assay as well as all following experiments. All *Salmonella* were grown in LB (Difco, BD, Thermo Scientific, Rockford, IL, USA) at 37°C, shaking at 220rpm for 14-16 hours prior to each use. Glycosyl hydrolase enzyme knockouts of *malS* and *nanH* were made as described previously [30]

### Human Cell Line and Growth Conditions

Human colonic carcinoma (Caco2) cell lines were obtained from ATCC (HTB-37) and grown as described previously [15]. Briefly, Caco2 cells were thawed from liquid nitrogen stocks stored in DMEM with 10% DMSO then grown in DMEM with 10 mM MOPS (Sigma, St. Louis, MO, USA), 10 mM TES (Sigma, St. Louis, MO, USA), 15 mM HEPES (Sigma, St. Louis, MO, USA), 2 mM NaH_2_PO_4_ (Sigma, St. Louis, MO, USA), 20% fetal bovine serum (HyClone Laboratories), 1% glutamax (Thermo Scientific, Rockford, IL, USA), 1% PenStrep (Thermo Scientific, Rockford, IL, USA) and 1% non-essential amino acids (Thermo Scientific, Rockford, IL, USA). Culture medium was renewed every three days. Caco2 cells were seeded at 10,000 cells/cm^2^ into 96-well plates then differentiated for 12 to 15 days for gentamicin protection assays.

### In Vitro Colonic Cell Infection Assays

Colonic cell infection assays were also performed according to Chen at al. [15] and Arabyan et al. [30], which adapted methods from Shah et al. [75, 76]. Oligosaccharides suspended at 1% (w/v) in serum-free DMEM were added to differentiated Caco2 cells and incubated for 15 mins. Following prebiotic pretreatment, stationary phase *S.* Typhimurium (n=3 biological replicates; multiplicity of infection=1000) was added to the pretreated Caco2 cells and incubated for 60 mins. PBS buffer (pH=7.2) was used to wash the cells and 50 mL Warnex buffer (AES Chemunex Canada, Inca, Montrealm QC, Canada) was applied to lyse the cells according to the manufacturer’s directions. Deactivation of Warner lysis was done with a 15 mins incubation at 95°C. Samples were then diluted 1:10 in nuclease-free water and stored at -20°C for qPCR quantification. Quantification of *S.* Typhimurium LT2 and 14028S cells was done via qPCR using primers previously validated by Arabyan et al [30], F: 5’-ACG CGG 313 TAT CAT CAA AGT GG - 3’; R: 5’ - ATC GGG TGG ATC AGG GTA AC - 3’. Significant differences in association across control and treatment were estimated using one-way ANOVA with Tukey test and graphed in GraphPad Prism V9 (GraphPad Software Inc, La Jolla, CA, USA).

### Metabolomics

Two different analytical setups for liquid chromatography coupled to mass spectrometry (LCMS) were used to survey a wider variety of non-volatile compounds. Samples were split between two capped LC vials, then were stored at -20°C prior to analysis. Non-volatile compounds from the culture supernatant were analyzed via LC-MS using a hydrophilic interaction chromatography (HILIC) column for hydrophilic and polar molecules and a reversed-phase (RP) column for nonpolar molecules according to the method by Borras et al. [77] and Aksenov et al. [78]. All samples were analyzed both via HILIC and RP on an Agilent 1290 series ultrahigh-performance LC system with an Agilent 6230 time-of-flight (TOF) mass spectrometer (Agilent Technologies, Santa Clara, CA, USA). An Agilent Jet Stream (AJS) nebulizer was used working in positive mode (+) and acquiring a mass range between 50 and 1700 Thomson (m/z) at 4 spectra/sec and high-resolution mode. Sheath gas temperature was 350°C, gas flow was 11 L/min, and fragmentor voltage was set at 120V. A model 6545 quadrupole TOF mass spectrometer (Agilent Technologies, Santa Clara, CA, USA) was used for final MS/MS compound identification. Water, acetonitrile, and 10% acetonitrile suspension mix were analyzed as blanks alongside the samples, which were all analyzed via injection of 5µL aliquots with samples housed in an autosampler maintained at 4°C. All LCMS analysis was performed with four biological replicates.

HILIC samples were analyzed on an Acuity UPLC bridged ethyl hybrid amide column (130Å, 1.7µm, 2.1mm × 100mm; Water, Milford, MA, USA). Water (A) and 90% acetonitrile in water (B) were used as mobile phases, both at pH 5 with ammonium acetate and acetic acid buffer. A linear gradient from 0% A to 10% A was applied post-injection in 20 mins with a flow rate of 0.3mL/min. The flow rate was then reduced to 0.2mL/min and phase A was increased to 95% in 10mins, for a total analysis time of 41mins. HILIC quality controls were a Water 1806006963 HILIC QC (Waters, Milford, MA, USA) and a custom-made QC. The custom QC consisted of 5µM carnitine, lysine, adenylputricine, aminocaproic acid, ornithine, tigonelline, alaninol, acetylcarnitine, 1-(2-pyramidyl)piperazine, methoxychalcone, cholecalciferol, 13-docosenamide and oleamide.

RP samples were analyzed on a Poroshell 120 EC-C18 column (2.7µm, 3.0mm x 50mm; Agilent Technologies, Wilmington, DE, USA) at 30°C. A mix of 1% phase A (60% acetonitrile in water) and 99% phase B (10% acetonitrile in isopropanol), both containing 10mM of ammonium formate and formic acid, was used as the initial mobile phase. The total analysis time was 24mins with a flow rate at 0.3mL/min, which consisted of phase B gradient reaching 30% post injection in 4mins, then rising to 48% B in 1min, 82% B in 17mins, and 99% B in 1min. The quality control was a standard solution Waters 6963 RP QC (Waters, Milford, MA, USA) and was injected along with the samples.

Agilent Mass Hunter Qualitative Analysis B.05.00SP1 software was used to examine the total ion LCMS chromatograms. The “Find by Molecular Feature” algorithm was used within a mass range from 50 to 1700Da for peak deconvolution. Molecular feature abundance was evaluated through integration of the extracted compound chromatograms (ECC) of the corresponding ions and then exported to .cef format. Mass Profiler Professional 12.1 software was used for peak alignment with a mass window of 40ppm, 26mDa, and a retention time shift of 0.5 and 1min for HILIC and RP, respectively. A peak table was made containing retention time in mins, molecular mass, and the intensity values (peak area) for each sample.

Resulting metabolic data was analyzed using MetaboAnalyst 5.0 [79]. Comparisons between treatments and across timepoints were made using both the statistical analysis [one factor] function and enrichment analysis modules. Samples were normalized by median, then all data was log_10_ transformed and scaled by mean-centering and standard deviation. Metabolite set enrichment analysis (MSEA) was performed using the KEGG database for reference.

### RNA Extraction

The 100K Pathogen Genome Project bacterial protocol [80] was used to extract *S*. Typhimurium RNA from the infection assays, while host cells were lysed by passaging cells through a 22-gauge needle [81]. Combined host and pathogen cells were pelted via centrifugation then suspended in Trizol LS Reagent (Cat #10296028, Thermo Fisher Scientific, Waltham, MA, USA). Host and pathogen RNA was extracted from the Trizol LS suspension following manufacturer’s instructions. The BioAnalyzer RNA kit (Agilent Technologies Inc., Santa Clara, CA, USA) and Nanodrop (Nanodrop Technologies, Wilmington, DE, USA) were used to confirm RNA purity (A_260/230_ and A_260/280_ ratios ≥ 1.8, ≤ 2.0) and integrity.

### RNAseq Library Preparation

RNAseq library preparation was performed exactly as outline in Chen et al. [15]. Briefly, the SuperScript Double-Stranded cDNA Synthesis kit (11917-010; Invitrogen, Carlsbad, CA, USA) was used to synthesize double-stranded cDNA following the manufacturer’s instructions. A NanoDrop 2000 spectrophotometer (Nanodrop Technologies, Wilmington, DE, USA) an Agilent 2100 BioAnalyzer (Agilent Technologies Inc., Santa Clara, CA, USA) was used to assay cDNA quality. All sequencing was done with 4 biological replicates in each condition.

The Kapa HyperPlus library preparation kit (kk814, KAPA Biosystems, Boston, MA, USA) with BIOO Scientific NEXTFlex adaptors (514105, BIOO, Austin, TX, USA) were used in the construction of the sequencing library. Library concentration was measured using the KAPA SYBR FAST qPCR kit Master Mix (2x) Universal (kk4903; KAPA Biosystems, Boston, MA, USA) on Bio-Rad CFX96 (Bio-Rad Laboratories, Hercules, CA, USA) and fragment size distribution was assessed using the High Sensitivity kit (Agilent Technologies Inc., Santa Clara, CA, USA). Libraries were indexed at eight libraries per lane and sequenced with PE150 on a HighSeq4000 at the California Institute for Quantitative Biosciences in the Vincent J Coates Genomics Sequencing Lab (Berkeley, CA, USA).

### Statistical Analysis for Differential Gene Expression

Sequence processing and analysis was done following the methods in Chen et al. [15]. Raw sequence reads were first trimmed with Trimmomatic [82] then aligned to the Ensembl GRCh38 human genome using HISAT2 [83] with an index downloaded on 07/16/22 (grch38_tran). Alignment was done in paired-end mode with soft clippings permitted and paired-end reads that did not map to the human genome were separated. Reads that did not align to the human genome were subsequently aligned using Bowtie2 [84] to the *Salmonella enterica* genome (GCA_003253385.1_ASM325228v1). Samtools [85] was used to compress alignment files from HISAT2 and Bowtie2 for output to differential expression analysis.

Differential expression analysis was performed in edgeR [86] from gene counts estimated by featureCounts in the Rsubread R package [87]. Human gene counts were produced using the Ensembl GRCh38.86.gtf annotation and *S. enterica* counts were generated from GCA_003253385.1_ASM325228v1.gtf annotation file. Human and *S.* Typhimurium gene count tables were entered separately into edgeR for normalization and differential expression. Genes with counts per million less than one and with expression in fewer than two samples per group were discarded and reads were normalized using the library size. Treatment groups contained pairwise comparisons, so the edgeR exact test was used for differential expression estimation. Significance was defined as adjusted *p*-value (FDR, this was done using a Bonferroni correction) of less than or equal to 0.05. No reads aligned to *Salmonella* from uninfected cells so no differential expression analysis was done for *Salmonella* reads in uninfected cells.

### Expression Data for Caco2 Cells

Qiagen’s Ingenuity Pathway Analysis software version 01-22-01 (IPA, Qiagen, Redwood City, CA, USA) was used to determine canonical pathways differentially expressed in Caco2 cells by prebiotic treatment. Canonical pathway mapping was performed in IPA and overlayed with experimental data. Expression data for TLRs was plotted using Prism 9 (GraphPad Software, La Jolla, CA, USA). Heatmaps were made using R Version 4.2.2 “Innocent and Trusting” along with the ComplexHeatmap Package version 2.16.0 [88]. All other plots related to expression data not from IPA were made using Prism 9 (GraphPad Software, La Jolla, CA, USA).

### Expression data for S. Typhimurium

*S.* Typhimurium differential expression data was uploaded to BioCyc SmartTables for further analysis (Pathway Tools version 27, SRI International, Menlo Park, CA, USA) [89]. Determination of enriched pathways was done using a two-tailed Fisher’s exact test (*p* ≤ 0.05) with the Pathway Tools *S.* Typhimurium strain 14028 genome. Enriched pathways from were plotted using Prism 9 (GraphPad Software, La Jolla, CA, USA) and heatmaps were made using R Version 4.2.2 “Innocent and Trusting” along with the ComplexHeatmap Package version 2.16.0 [88].

## Supplemental Figure titles

***Supplemental Figure 1.* Expression of virulence factors in *S.* Typhimurium and TLRs in Caco2 Cells**. (Left) Virulence factor expression in *S.* Typhimurium mixed with prebiotic treated Caco2 cells was measured as log_2_ fold change from *S.* Typhimurium exposed to untreated Caco2 cells. Orange represents *S.* Typhimurium add to the BioMos condition and purple represent the HMO condition *S.* Typhimurium. (Right) Toll-like receptor expression in prebiotic treated then infected Caco2 cells was evaluated by log_2_ fold change expression data with non-treated but infected Caco2 cells as control. Green represents BioMos treatment and purple represents HMO treatment.

***Supplemental Figure 2.* BioMos addition rescues invasion phenotype of glycosyl hydrolase knockouts.** Single gene knockouts of an amylase encoding gene (*malS*) and sialidase encoding gene (*nanH*) decreased the ability of S. Typhimurium LT2 to both adhere to and invade Caco2 cells. Pretreatment of the Caco2 cells with 1% BioMos for 60 mins restored the adhesion and invasion abilities of both the *malS* and *nanH* deletion as compared to a wild-type control. S. Typhimurium was co-incubated with the Caco2 cells for 60 mins and invasion and adhesion were measured using a gentamicin protection assay.

***Supplemental Figure 3.* Expression of a subset of transmembrane receptors in Caco2 cells diverges by prebiotic treatment.** Transmembrane receptor expression was measured in Caco2 cells pretreated with either HMO or BioMos then infected. Genes were clustered by Euclidian Distance and expression data is log_2_ fold change with non-prebiotic treated by infected Caco2 cells.

***Supplemental Figure 4.* Metabolic profiles of prebiotic treatments from infected Caco2 cells cluster by prebiotic type.** K-Means clustering, with cluster setting of 2, of metabolomes from *S.* Typhimurium infection across time and between prebiotic treatment groups across 60 min *S.* Typhimurium co-incubation. T_0_ STM v T_60_ STM compares metabolites from Caco2/STM treatment across T_0_ and T_60_. BioMos vs HMO compares metabolites across prebiotic pre-treatments at T_0_ and T_60_.

***Supplemental Figure 5.* Metabolic profiles of all conditions reveal distinct differences driven by both prebiotic treatment and *S.* Typhimurium presence/absence.** Correlation plot of untargeted metabolic profiles of all treatment combinations and individual replicates A-D for each combination created using MetaboAnalyst. Pink to red squares indicate positive correlation between metabolic profiles whereas gradients of blue represent negative correlations between profiles across sample type. Colored bars on the right-hand side of the plot indicate sample grouping by prebiotic type, pathogen presence/absence, and time course. Time 0 corresponds to 15 minutes post prebiotic addition and initiation of *S.* Typhimurium addition. Time 60 is 60 minutes post *S.* Typhimurium inclusion or 75 minutes after initial prebiotic addition when a *S.* Typhimurium was not added.

***Supplemental Figure 6.* Metabolites are significantly altered by treatment.** The metabolic profiles underlying the correlation plot in Supplemental Figure 4 were used to search for significance across all treatment combinations. The Kruskal Wallis Test in MetaboAnalyst revealed 252 significant metabolites out of 316 total.

***Supplemental Figure 7.* Graphical abstract of basic experimental set-up.** Caco2 cells (ATCC HTB-37) were grown and differentiated before being pre-treated with 1% HMO or BioMos for 15 mins. Stationary *S. enterica* sv. Typhimurium 14028 was then added to the cells and co-incubated for 60 mins. Cells and supernatant were collected, washed and stored for RNA-seq and metabolomics follow-up.

